# Draft genomes of clinical *Bacillus cereus* isolates AH725 and B06.009 closely related to reference strains ATCC 14579 and ATCC 10987

**DOI:** 10.1101/073924

**Authors:** Thomas H. A. Haverkamp, Nicolas J. Tourasse, Alexander J. Nederbragt, Kjetill S. Jakobsen, Anne-Brit Kolstø, Ole Andreas Økstad

## Abstract

*Bacillus cereus* strains B06.009 and AH725 were isolated from human cerebrospinal fluid and eye infection, respectively. Here we present their annotated draft genome sequences, which are highly similar to environmental *B. cereus* reference strains ATCC 10987 and ATCC 14579, underlining the opportunistic nature of *B. cereus* infections.

## Genome Announcement

The *B. cereus* group (*B. cereus sensu* lato) consists of at least seven approved species of Gram-positive rod-shaped spore-forming bacteria, including the human obligate pathogen *Bacillus anthracis*, entomopathogenic *Bacillus thuringiensis*, and the opportunistic human pathogen *B. cereus* (*B. cereus sensu stricto*). In genetic terms these bacteria could be characterized as one species (1), however species distinctions are kept mainly for reasons of variable pathogenicity patterns, thus largely being based on phenotypic traits that are plasmid-encoded (2).

Here, we present the draft genomes of *B. cereus* strains B06.009 and AH725. The B06.009 strain (Institut Pasteur, France) is to our knowledge the first *B. cereus* isolate from human cerebrospinal fluid infection to be sequenced. It belongs to *B. cereus* main population group III (3) and is identical in all seven MLST loci (ST2) to the *B. cereus* reference strain ATCC 10987, originally isolated in 1930 from contaminated cheese in Canada (4), employing the optimized Tourasse-Helgason MLST scheme (5, 6). *B. cereus* AH725 is an isolate originating from human eye infection (Skien Hospital, Norway; 1986) (7). It belongs to population group IV (ST33) and is identical in all seven Tourasse-Helgason MLST loci to the *B. cereus* type strain ATCC 14579, originally isolated from air in a cow-shed in the US in 1916. AH725 was previously shown to carry a 380 kb pXO1-like plasmid by pulsed-field gel electrophoresis (7).

*B. cereus* B06.009 and AH725 strains were grown to mid-log phase in LB medium at 30°C and shaking at 250 rpm. Cultures were pelleted and genomic DNA isolated using the Genomic DNA kit (Qiagen, Genomic-Tip 100/G), with a modified protocol employing treatment with 4% (24 mg) lysozyme and 200 U Mutanolysin prior to cell lysis.

Whole genome shotgun sequencing using 454-GS FLX was used to generate the draft genomes for AH725 and B06.009. The AH725 and B06.009 genomes were assembled using Newbler (version 1.1.03.24) into 655 and 520 contigs (>200bp), respectively. Annotation was performed using the NCBI prokaryotic genome annotation pipeline (version 2.10). The genomes have a length of 5,627,493 (AH725) and 5,198,273 (B06.009) bp and contain 5,763 and 5,237 predicted protein coding ORFs.

For B06.009 the *in silico* DNA-DNA hybridization (DDH) similarity to *B. cereus* ATCC 10987 was 99.70 % (http://ggdc.dsmz.de/distcalc2.php) (8, 9), and for AH725 when compared to the *B. cereus* type strain ATCC 14579, 90.30 % similarity (4). The Average Nucleotide Identity between B06.009 and ATCC 10987, as calculated with Jspecies (10), was 99.68%, and for AH725 and ATCC 14579 this was 98.47%. Thus, both DDH and ANI show a strikingly high level of genomic identity between clinical isolates AH725 or B06.009, and the reference genomes of environmental isolates isolated continents and decades apart, underlining the opportunistic nature of *B. cereus* infections.

The AH725 genome assembly does not contain the two prophages phBC6A51 (61Kbp) and phBC6A52 (38 Kbp), which are found in the reference strain ATCC14579 (2). Likewise, the reference strain ATCC10987 genome contains four predicted prophages (not annotated), one of which (position 393769-434007) is not identified in the B06.009 genome (4).

## Nucleotide sequence accession numbers

Both whole genome shotgun projects have been deposited in GenBank, under the accession numbers LEBX00000000 (AH725) and LEBZ00000000 (B06.009). The versions described in this paper are the first versions.

## Acknowledgments

We thank the Norwegian Sequencing Centre (NSC), in particular Ave Tooming-Klunderud, for 454 sequencing.

